# Assessment of biological role and insight into druggability of the *Plasmodium falciparum* protease plasmepsin V

**DOI:** 10.1101/426486

**Authors:** Alexander J. Polino, S. Nasamu Armiyaw, Jacquin C. Niles, Daniel E. Goldberg

**Affiliations:** Division of Infectious Diseases, Department of Medicine, Washington University School of Medicine, St. Louis, Missouri, USA.; Department of Biological Engineering, Massachusetts Institute of Technology, Cambridge, MA, USA.

## Abstract

Upon infection of a red blood cell (RBC), the malaria parasite *Plasmodium falciparum* drastically remodels its host by exporting hundreds of proteins into the RBC cytosol. This program of protein export is essential for parasite survival, hence there is interest in export-related proteins as potential drug targets. One proposed target is plasmepsin V (PMV), an aspartic protease that cleaves export-destined proteins in the parasite ER at a motif called the *Plasmodium* export element (PEXEL). This cleavage is essential for effector export across the vacuolar membrane. Despite long-standing interest in PMV, functional studies have been hindered by the failure of current technologies to produce a regulatable lethal depletion of PMV. To overcome this technical barrier, we designed a facile system for stringent post-transcriptional regulation, allowing a tightly controlled, tunable knockdown of PMV. Under maximal knockdown conditions, parasite growth was arrested, validating PMV as essential for parasite survival in RBCs. We found that PMV levels had to be dramatically depleted to affect parasite growth, suggesting that the parasite maintains this enzyme in substantial excess. This has important implications for antimalarial development. Additionally, we found that PMV-depleted parasites arrest immediately after invasion of the host cell, suggesting that PMV has an unappreciated role in early development that is distinct from its previously reported role in protein export in later-stage parasites.

**Importance:** Malaria is endemic to large swaths of the developing world, causing nearly 500,000 deaths each year. While infection can be treated with antimalarial drugs, resistance continues to emerge to frontline antimalarials, spurring calls for new drugs and targets to feed the drug development pipeline. One proposed target is the aspartic protease plasmepsin V (PMV) that processes exported proteins, enabling the export program that remodels the host cell. This work uses facile genetic tools to produce lethal depletion of PMV, validating it as a drug target and showing that PMV is made in substantial excess in blood-stage parasites. Unexpectedly, PMV depletion leads to parasite death immediately after invasion of RBCs, distinct from other disruptions of the export pathway. This suggests that PMV inhibitors could lead to relatively rapid parasite death, and that PMV has additional unexplored role(s) during RBC infection.

## Introduction

Malaria remains a scourge of the developing world, causing nearly 500,000 deaths per year, with the overwhelming majority due to infection by the intracellular parasite *Plasmodium falciparum* (1). While the life cycle of *P. falciparum* includes replication in both the liver and blood, symptomatic human disease is caused by infection of red blood cells (RBCs) (2). Upon infection of a host RBC, the parasite executes a dramatic protein export program, sending hundreds of parasite proteins through the secretory system and across the surrounding vacuole (parasitophorous vacuole, PV), into the host cytosol (3, 4). These exported effectors drastically remodel the host cell, setting up new solute permeability pathways, modifying the RBC shape and rigidity, and reconstituting trafficking machinery in the RBC cytosol to send parasite-encoded adhesins to the RBC surface (3, 4). These adhesins mediate binding of infected RBCs to vascular endothelia allowing parasites to avoid splenic clearance. Adherent parasites in the brain can cause vascular blockage leading to death in severe cases (2).

Due to the central role of protein export in the survival and virulence of *P. falciparum*, there has been considerable interest in this process as a source of drug targets. One target of interest has been the parasite aspartic protease plasmepsin V (PMV). PMV processes exported proteins in the parasite ER by cleaving them co-translationally in a variant signal recognition particle complex (5) at a conserved amino acid export motif termed the *Plasmodium* export element (PEXEL) (6–8). PMV is highly specific for RxL in the PEXEL and cleaves after the leucine (9–12). PEXEL processing is a critical step in protein export, as mutations in PEXEL that block PMV processing also block protein export (9, 10). Furthermore, processing of PEXEL proteins is likely an essential function in the parasite, as PMV knockouts have thus far been unobtainable in the rodent malaria parasite *P. berghei* (6, 7, 13), a catalytic aspartate mutant of PMV acts as a dominant negative, reducing growth of cultured *P. falciparum* (7), and a conditional PMV knockout in *P. falciparum* is lethal (14).

In recent years, a number of tools have been used to study PMV, including peptidomimetic inhibitors that block its function *in vitro* and appear to be on-target and lethal to parasites in culture (15–17), as well as a crystal structure of the *P. vivax* PMV (18). However, study of PMV function has been hindered due to an inability of previous knockdown methods to yield a phenotype in RBC culture. The most robust knockdown described to date used the *glmS* ribozyme system to reduce PMV levels 10-fold. However, this had no measurable effect on parasite growth (15). Here, we sought to apply the recently described TetR-DOZI aptamer system for stringent and tunable regulation of PMV (19). This system has been shown to deplete a reporter gene 45 to 70-fold when aptamers are installed in both the 5’ and 3’ untranslated regions (UTRs) of the target gene (19). However, cloning such a construct in traditional plasmid systems requires the assembly and maintenance of large circular plasmids that are prone to deletions and vector rearrangements during propagation in *E. coli* (20). To overcome this technical challenge, we assembled a number of tools for genetic manipulation onto a previously described linear vector. Using this new vector system, we achieved substantially greater depletion of PMV than was previously described. We found that depletion of PMV was lethal to parasites, validating PMV as an essential protein in RBC culture. By tuning the degree of knockdown, we showed that PMV is maintained at substantial excess during RBC infection and must be suppressed to nearly undetectable levels to affect parasite growth. Finally, we found that PMV-depleted parasites die immediately after invasion in a manner distinct from that of disruptions of other protein export machinery, suggesting PMV may have additional roles beyond those previously described.

## Results

### Constructions of a linear vector for aptamer knockdowns

To overcome the challenges associated with maintaining *P. falciparum* genomic material in circular plasmids, we utilized the pJAZZ linear vector system (20) as a chassis for DNA assembly operations. This system has previously been used to manipulate large [A+T]-rich genomic fragments, including those derived from the rodent malaria parasite, *P. berghei* (20, 21). We constructed a plasmid (pSN054) to allow facile cloning, endogenous tagging, robust regulation of expression and/or inducible knockout of *P. falciparum* genes (Fig. 1A, Fig. S1). pSN054 has the following features: a single 5’ aptamer, a 10x array of 3’ aptamers, a fixed regulatory protein (TetR-DOZI repressor complex)(19), a parasitemia tracking component (Renilla luciferase), a drug selection marker, a cassette for generation of sgRNA for CRISPR/Cas9 genome editing, modular cloning sites for inserting left homologous regions (LHRs) and right homologous regions (RHRs) for homologous repair, modularized affinity tags for tagging genes of interest at the N-terminus or C-terminus and loxP sites for gene excision if needed (Fig. 1).

**Figure 1.**
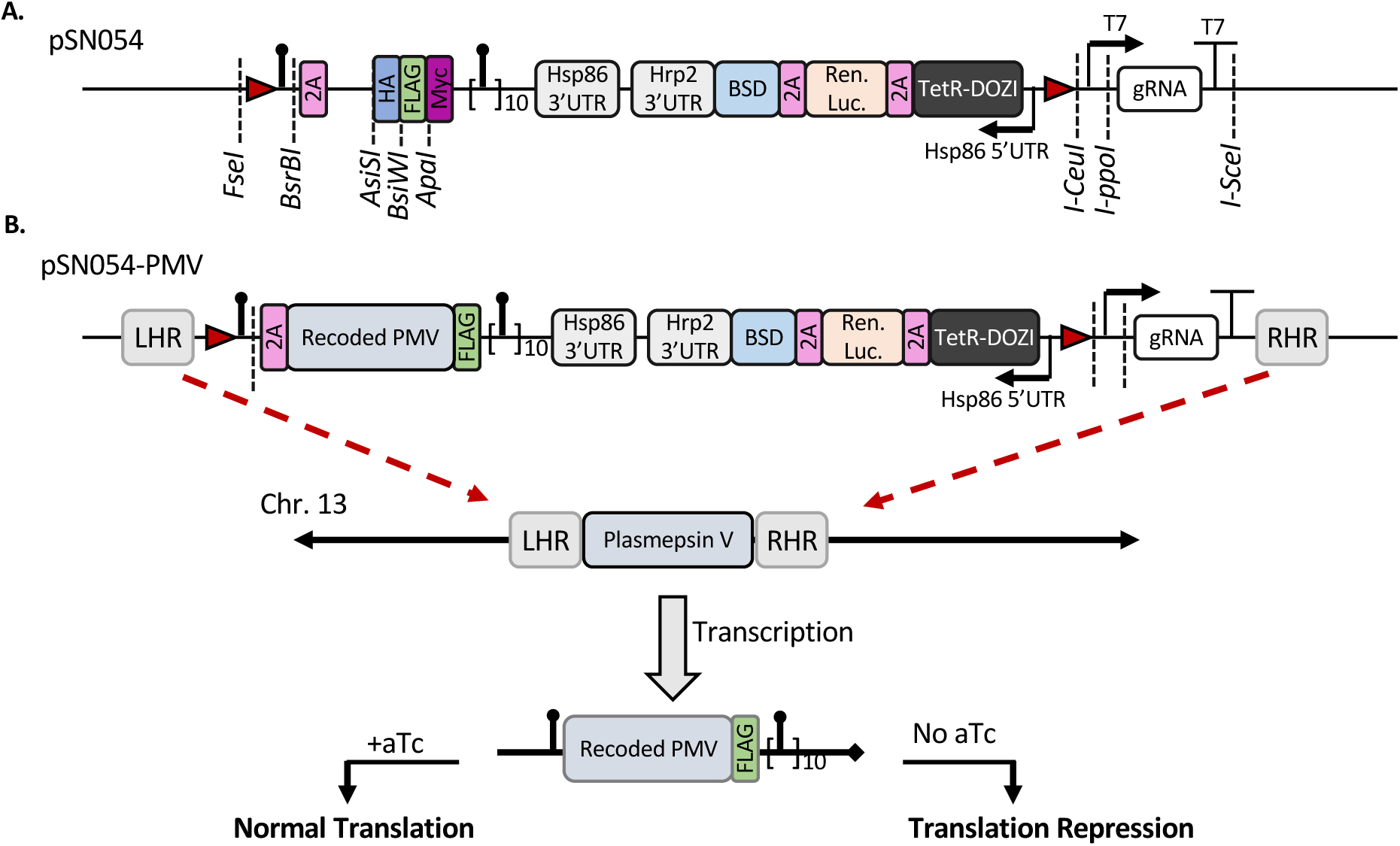
Architecture of pSN054 and application to editing PMV. (A) Schematic of pSN054 showing restriction sites for cloning (dashed lines), loxP sites for DiCre operations (red triangles), and aptamers (black lollipops). 2A (pink) is a viral skip peptide. Restriction sites allow the choice of N- or C-terminal 3xHA (blue), FLAG (green), or Myc (purple). The plasmid also contains an expression cassette driving production of the Tet repressor and DOZI helicase fusion (TetR-DOZI, black), Renilla luciferase (Ren. Luc.), and blasticidin-S deaminase selectable marker (BSD). The T7 expression cassette is for driving transcription of CRISPR guide RNAs (gRNA), which can be inserted by cutting with I-ppoI or AflII (contained within the I-ppoI cut site). (B) Diagram showing the cloning strategy for editing of the PMV locus. Left and right homologous regions (LHR and RHR) were inserted at FseI and I-SceI respectively, while the recoded gene sequence was inserted into plasmid cut with AsiSI and BsiWI. The endogenous sequence of PMV was disrupted by CRISPR/Cas9 gene editing. When transcribed, aptamers are bound by TetR-DOZI in the absence of aTc, repressing translation. In the presence of aTc, TetR-DOZI does not bind the aptamers and translation occurs as normal.

To utilize pSN054 for genome editing, a gene of interest must be recodonized to replace the current gene, preventing aberrant homologous repair from truncating the construct. To facilitate repair, 400-600bp of left and right homologous regions (LHR and RHR), corresponding to 5’UTR and 3’UTR of a gene, are cloned into the FseI and I-SceI restriction sites respectively (Fig. S1). This intermediate vector can be used to create gene knockouts since it does not possess the coding sequence of the gene of interest. A subsequent Gibson assembly is then performed to insert the coding sequence of the gene. There are several choices for this step depending on the tag one desires and whether this tag is N or C-terminal. pSN054 contains the 2A skip peptide (23) such that cloning into the AsiSI site produces a protein with no tag on the N-terminus. If an N- or C-terminal tag is desired, the relevant cloning site is used for gene insertion (Fig. 1A). This donor plasmid can be adapted for use with the T7-RNAP CRISPR/Cas9 system (24) by cloning an sgRNA into the AflII site. Alternatively, a U6 promoter-containing gRNA plasmid (25) can be used and co-transfected with a finished pSN054 donor plasmid.

### TetR-DOZI aptamer system enables lethal knockdown of PMV

To apply pSN054 to PMV, we cloned pieces of the 5’UTR and 3’UTR into FseI and I-SceI respectively, as well as a recoded version of the PMV coding sequence into plasmid cut with AsiSI and BsiWI (Fig. 1B). This plasmid enables us to simultaneously tag PMV with a C-terminal FLAG tag and regulate expression with the TetR-DOZI system (Fig. 1B). The construct was transfected into NF54 (26), and parasites selected and cloned. Incorporation of the construct into the genome was verified by Southern blot (Fig. 2A). Expression of a FLAG-tagged protein of the expected size was verified by western blot (Fig. 2B). Similarly, a western blot with anti-PMV verified that modification of this locus did not substantially change PMV expression levels of the edited line relative to the parent (Fig. 2C).

**Figure 2.**
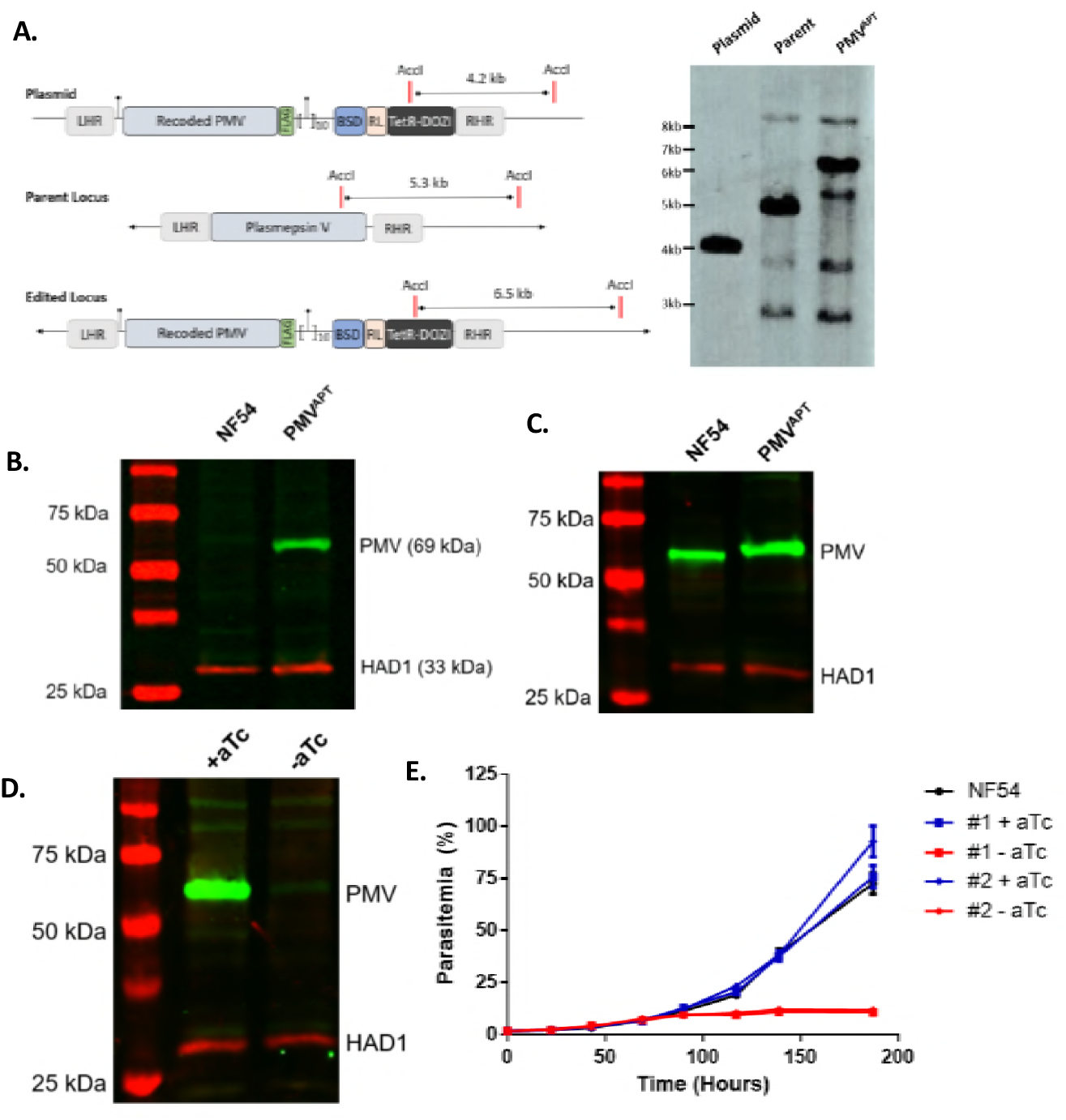
pSN054 enables lethal knockdown of PMV. (A) Integration at the proper locus was verified by Southern blot, with the right homologous region (RHR) used as a probe. (B) Protein tagging was verified by western blot using an anti-FLAG antibody as well as an antibody to a cytosolic sugar phosphatase HAD1 as a loading control. (C) Expression levels were compared to the parent strain by western blot using anti-PMV and anti-HAD1 antibodies. (D) Knockdown of PMV was initiated by washing aTc from ring-stage cultures and adding either 500nM aTc (+aTc) or an equal volume of DMSO (-aTc). After 72 hours, knockdown was assessed by western blot, probing with anti-PMV and anti-HAD1. (E) Parasites were split +/- aTc as above and growth monitored by flow cytometry daily. Experiment was done twice in technical triplicate. A representative experiment is shown, with data points representing mean and error bars standard deviation. Uncut gels are shown in Fig. S2.

We then utilized the TetR-DOZI aptamer system to post-transcriptionally deplete PMV. Knock down was initiated by washing out anhydrotetracycline (aTc) from young ring-stage parasites in RBC culture. In the absence of aTc, PMV levels were depleted about 50-fold (Fig. 2D) and parasite growth was arrested after 96 hours (Fig. 2E). This confirms that PMV is essential for intraerythrocytic growth and showcases the ability of the TetR-DOZI aptamer system to drive more substantial depletion of proteins than previously possible.

### PMV is maintained in substantial excess during infection

We next sought to utilize the tunability of the TetR system to determine the amount of PMV required for parasite survival. To this end, we titrated aTc levels and followed parasite growth by flow cytometry (Fig. 3A). Parasites maintained in 3 nM aTc or above grew normally. Parasites maintained in DMSO or 1 nM aTc arrested by 96h, while parasites maintained in 2 nM aTc survived an additional cycle before arresting at 120h (Fig. 3A). Given this, we sought to determine the amount of PMV depletion necessary to affect parasite growth. We synchronized parasites and washed out aTc from cultures containing predominantly young ring-stage parasites, then harvested samples for western blot at 72h following washout to compare the PMV levels in parasites that would go on to survive another cycle (2 nM aTc) to those that would arrest in the next 24h (1 nM aTc). Over three independent experiments, we found that 8% of wildtype PMV (2 nM aTc) was sufficient to support another cycle of growth, while 3% (1 nM aTc) was insufficient and led to death upon reinvasion (Fig. 3B, C). This suggests that PMV is maintained in enormous functional excess in parasite culture.

**Figure 3.**
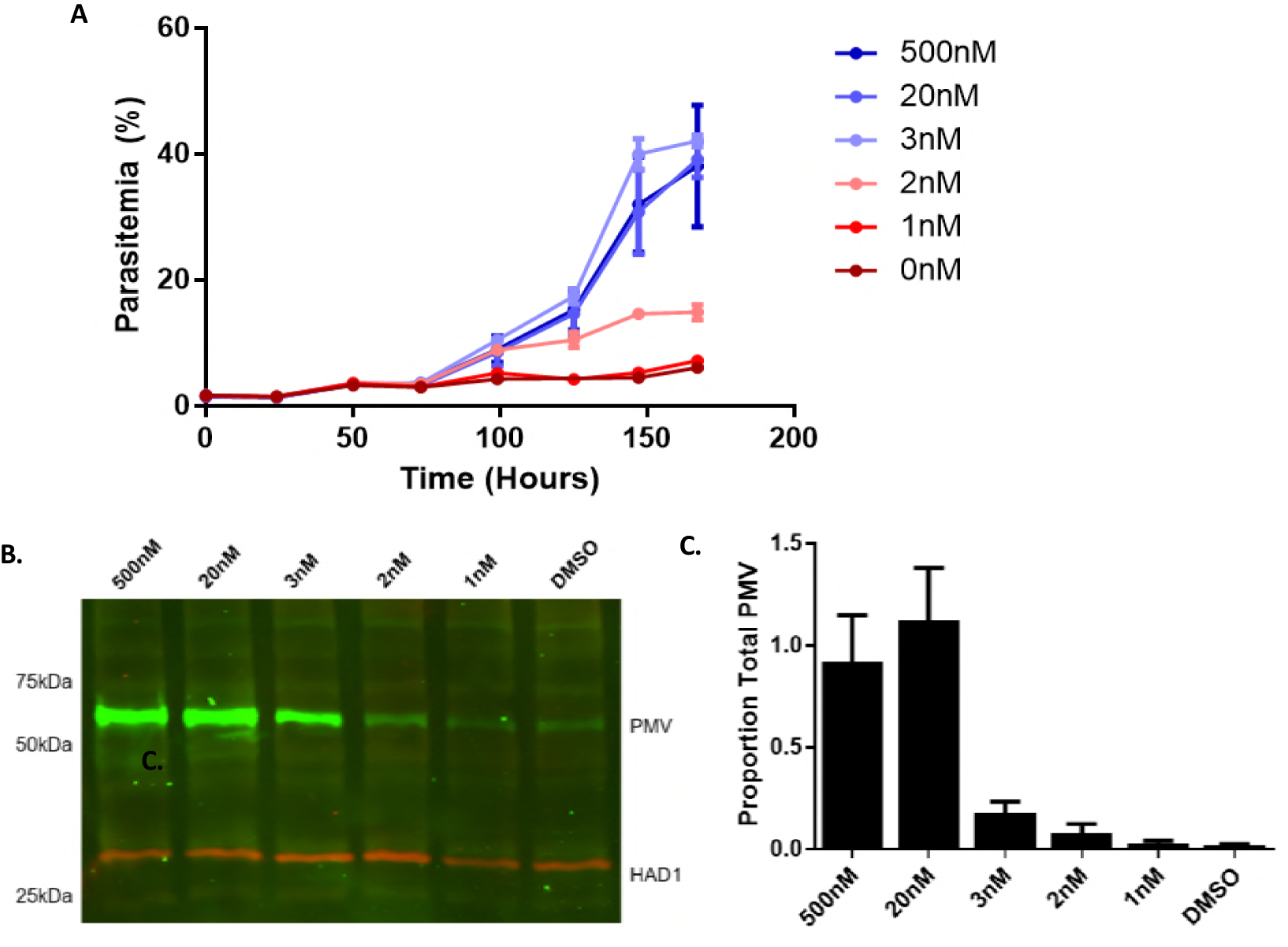
Tunable regulation shows PMV is maintained in substantial excess. (A) aTc was washed from parasite cultures as above, and parasites maintained in 500nM, 20nM, 3nM, 2nM, 1nM, or 0nM (DMSO) aTc. Growth was monitored by flow cytometry and data represented as in Fig. 2. Three experiments were performed with each sample done in triplicate. A representative experiment is shown. (B) Samples were prepared as in (A) but harvested at 72h for western blot. Three experiments were performed. A representative blot is shown in (B), and quantification of the three experiments is shown in (C). Bars represent the mean of PMV signal over HAD1 signal over the three experiments; error bars represent the SEM. Uncut gels are shown in Fig. S3.

### PMV depleted parasites arrest early in life cycle

Given the canonical role of PMV in parasite protein export, we expected PMV depletion to phenocopy disruption of other critical export machinery such as components of the *Plasmodium* translocon for exported proteins (PTEX) which mediates translocation of effectors across the PV membrane (3, 27, 28). PTEX components Hsp101, PTEX150 and EXP2 are all essential in parasite culture, and previously described depletion of any of these caused parasite arrest during the early trophozoite stage (29–31). Therefore, we were surprised to find that PMV depletion caused growth arrest very early in the intraerythrocytic development cycle, shortly after invasion (Fig. 4A). Arrested parasites were further investigated by transmission electron microscopy and were found to have gross structural abnormalities (Fig. 4B). PMV-depleted parasites generally showed more electron density throughout the parasite cytosol, failure to expand much beyond the size of merozoites, and large unidentified vacuolar structures within the parasite. In contrast, when parasites were fixed as schizonts from the preceding cycle, there was no obvious morphological defect (Fig. 4C). This suggests that PMV plays some critical role(s) in parasites immediately after invasion, distinct from its canonical protein export role.

**Figure 4.**
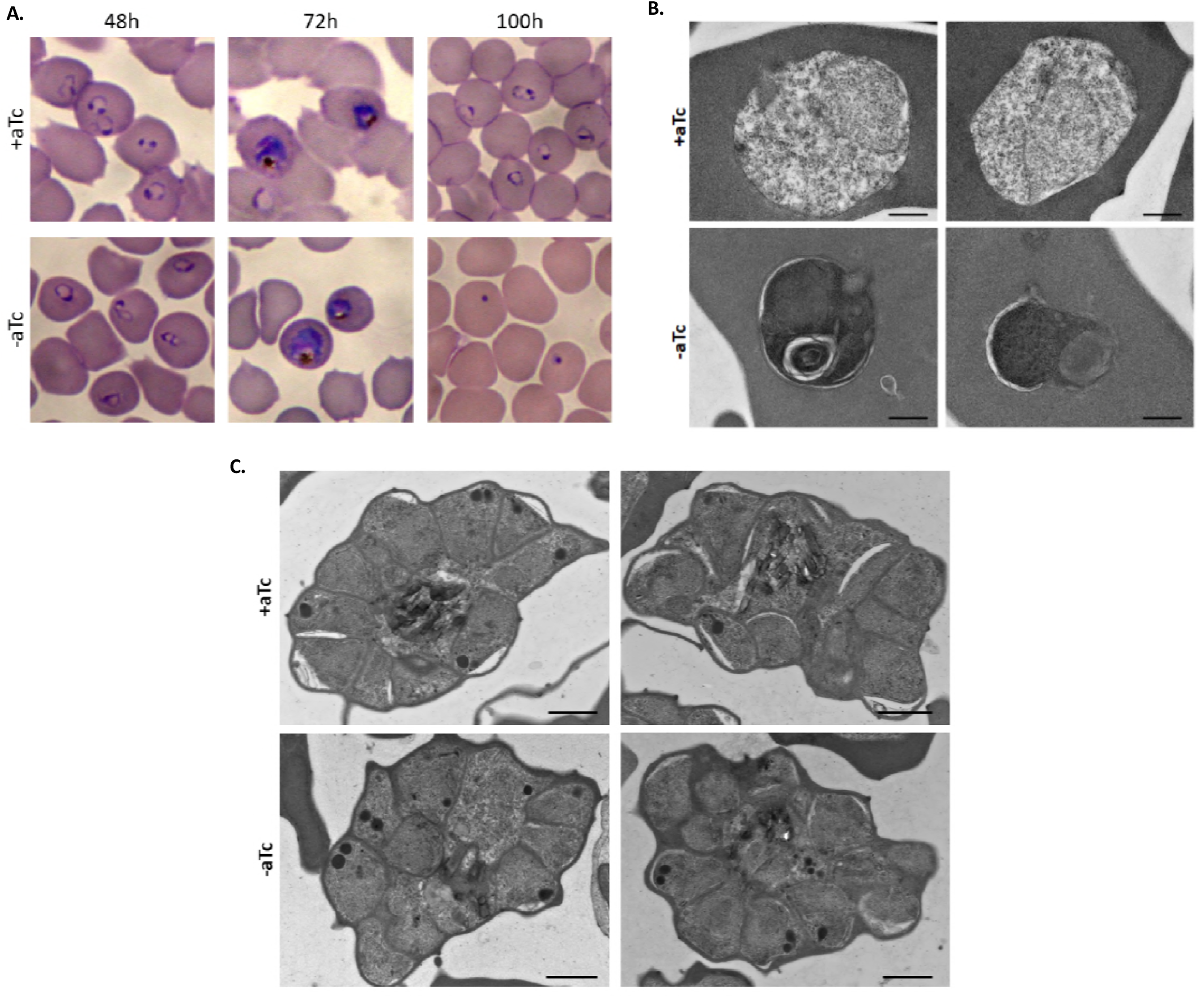
PMV-depleted parasites arrest early in life cycle. (A) . (D) aTc was washed from ring-stage parasites, which were then maintained in 500nM aTc (+aTc) or an equal volume of DMSO (-aTc). Parasite growth was monitored by Hemacolor-stained thin smear. (B) Parasites maintained as above. At 96h, parasites were fixed and early ring-stage parasites visualized by transmission electron microscopy. (C) Parasites were maintained as in (B) except schizonts were harvested at 90h following aTc washout. Scale bars represent 1 μm.

### PMV inhibitors kill parasites at multiple times during the intraerythrocytic development cycle

Previous work indicated that PMV inhibitors were lethal to the parasite only in early trophozoites, consistent with depletion of PTEX components (18). Since this contradicts our finding that PMV depletion leads to parasite growth arrest early after invasion, we sought to recapitulate the previously described early trophozoite death as well as our early post-invasion death with the peptidomimetic PMV inhibitor WEHI-842 (16, 18). We treated synchronized ring-stage parasites or schizonts with 5 μM WEHI-842 for an 8-hour window, then monitored parasites by thin smear. We found that ring-stage parasites treated with WEHI-842 arrested as early-trophozoites as previously described (Fig. 5A) However, parasites treated with WEHI-842 beginning in schizogony arrested immediately after invasion, as was seen in our genetic PMV depletion line (Fig. 5B). Taken together, our data suggest that parasites are sensitive to PMV inhibition during two points in the life cycle. The first is immediately after invasion. The second is in early trophozoites and is phenocopied by PTEX disruptions.

**Figure 5.**
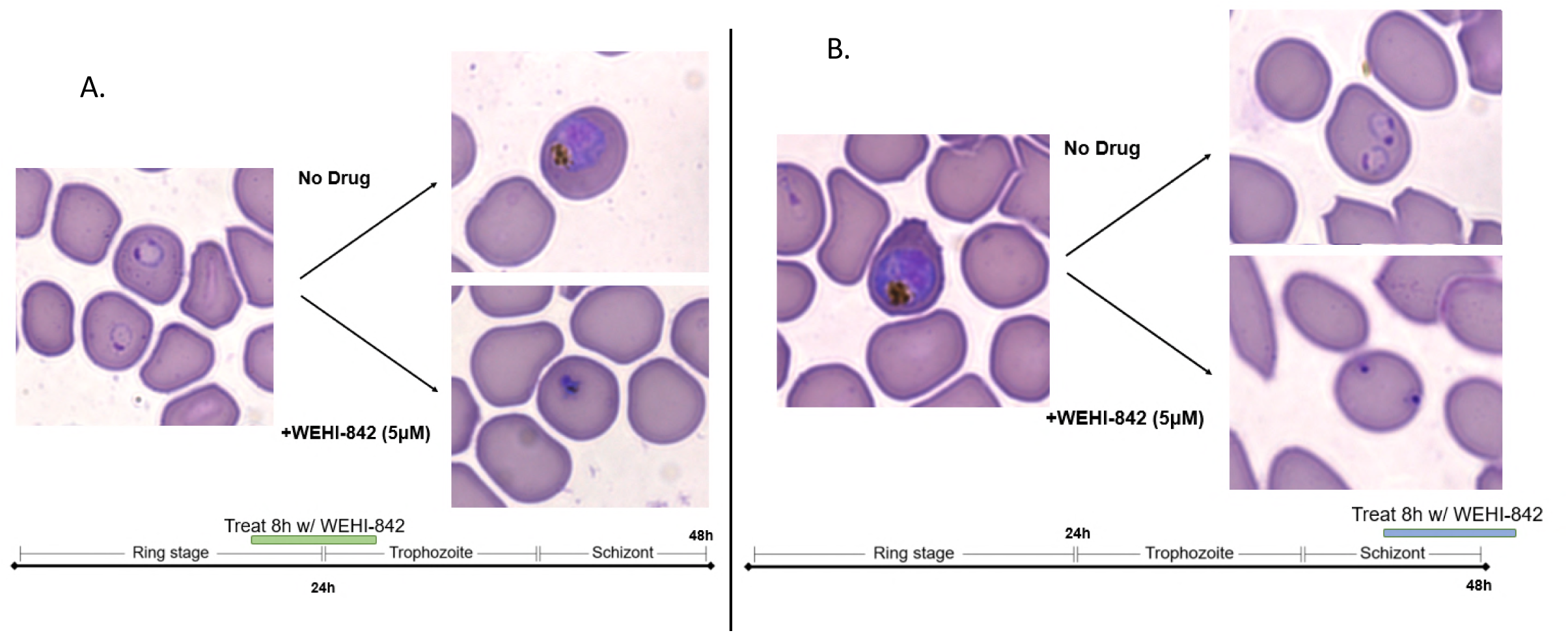
PMV inhibition arrests parasite growth at two distinct points in the life cycle. Parasites were synchronized to within 3 hours, then treated with 5 μM WEHI-842 for an 8-hour window beginning either in (A) ring-stage or (B) schizont. At the beginning and end of each window, parasites were monitored by Hemacolor-stained thin smear. Experiment was performed twice; representative images from one experiment are shown.

## Discussion

We report the first regulatable knockdown that lowers PMV levels enough to provide a lethal phenotype. This work overcomes the technical limitations of past knockdown systems by utilizing the TetR system with aptamers on both the 5’ and 3’ end of the gene of interest. This manipulation was greatly facilitated by the plasmid described here, pSN054, which enables a suite of previously described genetic tools to be utilized with relative ease for gene editing to achieve protein tagging, regulation of gene expression, parasite growth monitoring and inducible knockout as required.

An unexpected result is that PMV depletion does not phenocopy the disruption of PTEX components (29–31). The fact that the post-invasion death phenotype described here is not seen in disruption of other export machinery suggests that PMV has some role independent of protein export that is essential for parasite survival in RBCs. A reasonable hypothesis is that this role could be the cleavage of a critical substrate early in the life cycle to allow it to perform an essential activity. In this case, the substrate would likely be acting within the parasite or within its vacuole, since disruption of Hsp101 function with a destabilization domain blocked nearly all exported effectors within the vacuole but only arrested growth in early trophozoites (29). The identity of this substrate is an interesting avenue for future research. Alternatively, early death could be a non-specific result of PMV deficiency, such as a buildup of uncleaved PEXEL proteins in the ER. Consistent with this, treatment with the canonical ER-stress inducer DTT arrested growth in *P. falciparum* with similar morphology by Giemsa stain to that caused by PMV depletion described here (32).

One encouraging note for the development of PMV inhibitors as antimalarials is our finding that PMV inhibition can lead to parasite death at two distinct points within intraerythrocytic development. Knockdown of PTEX components seem to cause growth arrest only at one point in blood-stage parasites (29–31). Specifically, when Hsp101 or EXP2 protein levels were reduced to barely detectable levels in late-stage parasites, the parasites proceeded to invade and develop normally before arresting at the early trophozoite stage (30-31). Due to this, drugs inhibiting the function of PTEX components may take up to a full intraerythrocytic cycle (48 hours) to reach the point in the cycle where growth arrest occurs. In contrast, PMV inhibitors may be able to arrest growth more quickly by acting upon intraerythrocytic parasites at more than one point in the life cycle. This may bring a potential PMV inhibitor in line with recommendations that an ideal antimalarial act within 24 hours (33). However, this beneficial characteristic is counterbalanced by the important insight from our study that PMV must be suppressed to barely detectable levels to affect parasite growth. This may make PMV a challenging target for drug development as concentrations of inhibitors far beyond the IC_50_ calculated with purified enzyme may be required to cause parasite death. Peptidomimetic inhibitors of PMV that have been developed are generally greater than 100-fold less potent in culture than on isolated enzyme (15, 17, 18). It has been presumed that potency against parasites is limited by cellular permeability. Our functional genetics data would suggest that an additional, and possibly major, component of the potency drop-off is the need to inhibit nearly all the cellular enzyme to kill parasites. We usually measure enzyme inhibition in terms of an IC_50_ value, but in this case an IC_99_ might be more relevant to whole parasite killing.

Taken together, our study provides further data on the proposed antimalarial drug target plasmepsin V. Future work is needed to determine if PMV is maintained at excessive levels *in vivo* as it is *in vitro*, and to elucidate the cause of growth arrest after invasion in PMV-depleted parasites.

## Materials and Methods

### Parasite lines and culture

*P. falciparum* strain NF54^attB^ (referred to as NF54 throughout) was used as a parent strain for transfections (26). Asexual parasites were cultured in RPMI 1640 (Gibco) supplemented with 0.25% (w/v) Albumax, 15mg/L hypoxanthine, 110mg/L sodium pyruvate, 1.19g/L HEPES, 2.52g/L sodium bicarbonate, 2g/L glucose, and 10mg/L gentamycin. Deidentified RBCs were obtained from the Barnes-Jewish Hospital blood bank (St. Louis, MO).

### Generation of knockdown line

The construct for aptamer regulation of PMV was constructed using pSN054, described above. The right homologous region (3’ UTR) was amplified from NF54 genomic DNA using primers AGTGGTGTACGGTACAAACCCGGAATTCGAGCTCGGGGAATCAACATAGAAACGTTAAAG and GATTGGGTATTAGACCTAGGGATAACAGGGTAATGTACTAGGTCATTTTCTTTATTTTAC, and cloned into the I-SceI site using Gibson Assembly (NEB). The left homologous region (5’ UTR) was amplified from NF54 genomic DNA using primers TTGGTTTTCAAACTTCATTGACTGTGCCGACATTAATTTGTGTAACATATAAATATGTAG and AAGTTATGAGCTCCGGCAAATGACAAGGGCCGGCCCTTTCCTTAAAAAATAATTATTGAT, and cloned into the FseI site. PMV was codon-optimized for expression in *Saccharomyces cerevisiae* and synthesized as gene blocks by Integrated DNA Technologies (Coralville, IA) then cloned into the vector at the AsiSI site. The plasmid was grown in BigEasy Electrocompetent Cells (Lucigen) with 12.5 μg/mL chloramphenicol and 0.01% (w/v) arabinose.

CRISPR/Cas9 editing was performed as previously described (25). Guide RNA sequences were inserted into the pAIO vector by annealing oligonucleotides of the sequences ATTAAGTATATAATATT**TGTAATGGTTGTAAAGATTG**GTTTTAGAGCTAGA and TCTAGCTCTAAAAC**CAATCTTTACAACCATTACA**AATATTATATACTTAAT and inserting them into BtgZI-cut pAIO by In-Fusion HD Cloning (Clontech). pAIO was maintained in XL10 Gold cells (Agilent Technologies). Bold sequences represent the gRNA site.

For each transfection, 100 μg of donor vector and 50 μg of pAIO were transfected into early ring-stage parasites in 2mm gap electroporation cuvettes (Fisher) using a BioRad Gene Pulser II. Transfectants were maintained in 0.5 μM anhydrotetracycline (aTc; Cayman Chemical) and were selected beginning 24 hours post transfection with Blasticidin S (2.5 μg/mL; Fisher). Parasites were obtained from several independent transfections and clones obtained by limiting dilution. **Validation of PMV^APT^ line.** Proper integration of our construct was verified by Southern Blot as in (24). For a probe, the right homologous region was amplified from NF54 genomic DNA using primers described above.

To verify tagging of protein, schizonts of NF54 and PMV^APT^ were first synchronized by purifying on magnetic columns (Miltenyi Biotech) then allowed to invade fresh uninfected RBCs for 3 hours before remaining schizonts were cleared with 5% sorbitol. Parasites were then allowed to progress for 40h, then RBCs lysed with cold PBS + 0.035% saponin. Lysates were centrifuged to pellet parasites and remove excess hemoglobin, then parasites were lysed in RIPA (50mM Tris, pH 7.4; 150mM NaCl; 0.1% SDS; 1% Triton X-100; 0.5% DOC) plus HALT-Protease Inhibitor Cocktail, EDTA-free (Thermo Fisher). Lysates were centrifuged at high speed to pellet and remove hemozoin. Cleared lysates were then diluted in SDS sample buffer (10% SDS, 0.5M DTT, 2.5mg/mL bromophenol blue, 30% 1M Tris pH 6.8, 50% glycerol) and boiled. Lysates were separated by SDS-PAGE, then transferred to 0.45 μm nitrocellulose membrane (BioRad). Membranes were blocked in PBS + 3% bovine serum albumin, then probed with primary antibodies mouse anti-PMV 1:25 (34) or anti-FLAG 1:500 (M2, Sigma), and rabbit anti-HAD1 1:1000 (35). Membranes were washed in PBS + 0.1% Tween 20, then incubated with secondary antibodies goat anti-mouse IRDye 800CW 1:10,000 (Licor) and donkey anti-rabbit IRDye 680RD 1:10,000 (Licor). Membranes were then washed in PBS + 0.1% Tween 20 and imaged on a Licor Odyssey platform.

### Assessment of knockdown

To assess the effect of PMV knockdown on parasite growth, aTc was removed from cultures by washing 3 times for 5 minutes each in media without aTc, then either 500nM aTc (“+ aTc”) or DMSO (“- aTc”) was added and growth followed by flow cytometry. Flow cytometry data is plotted with each point representing the mean of three technical replicates with error bars showing the standard deviation. Experiments were done three times unless otherwise noted; a representative experiment is shown.

For titrations of aTc, parasite cultures were washed as above to remove aTc, then cultures were maintained in media containing the shown concentrations of aTc or DMSO (“0 nM aTc”). To determine the effect of aTc concentration on PMV levels, cultures were prepared as above and samples taken for western blot 72 hours after aTc was removed. Sample preparation and western blotting was done as above (see **Validation of PMV^APT^ line**). To quantitate PMV titration blots, the 500nM sample was diluted out by factors of two to draw a standard curve correlating PMV signal to relative amount of the 500nM sample (see Figure S3). Blots were quantitated using Licor Image Studio. The experiment was performed three times and the mean for those three experiments is plotted with error bars representing the standard error of the mean.

### WEHI-842 treatment

Parasites were synchronized to within 3 hours as above (see **Validation of PMV^APT^ line** above, paragraph 2). Then either 5 μM WEHI-842, or an equal volume of DMSO was added to late rings (15h after invasion initiated; parasites 12-15h old), or late trophozoites (44 hours after invasion initiated; parasites 41-44h old). Parasites were incubated with drug or DMSO for 8h, then assessed by thin smear. The experiment was performed twice, with similar results in each. Representative images from one experiment are shown.

### Microscopy

Parasites monitored by thin smear were dyed using Harleco Hemacolor stains (MilliporeSigma). Images were taken using a Zeiss Axio Observer. D1 at the Washington University Molecular Microbiology Imaging Facility. For transmission electron microscopy, infected RBCs were fixed in 2% paraformaldehyde/2.5% glutaraldehyde (Polysciences Inc., Warrington, PA) in 100 mM sodium cacodylate buffer, pH 7.2 for 1 hr at room temperature. Samples were washed in sodium cacodylate buffer at room temperature and postfixed in 1% osmium tetroxide (Polysciences Inc.) for 1 hr. Samples were then rinsed extensively in dH_2_0 prior to en bloc staining with 1% aqueous uranyl acetate (Ted Pella Inc., Redding, CA) for 1 hr. Following several rinses in dH_2_0, samples were dehydrated in a graded series of ethanol and embedded in Eponate 12 resin (Ted Pella Inc.). Sections of 95 nm were cut with a Leica Ultracut UCT ultramicrotome (Leica Microsystems Inc., Bannockburn, IL), stained with uranyl acetate and lead citrate, and viewed on a JEOL 1200 EX transmission electron microscope (JEOL USA Inc., Peabody, MA) equipped with an AMT 8-megapixel digital camera and AMT Image Capture Engine V602 software (Advanced Microscopy Techniques, Woburn, MA).

### Database links

Sequences for genes used in this study were obtained from PlasmoDB (Release 38) using the following gene IDs: plasmepsin V (PF3D7_1323500), HAD1 (PF3D7_1033400). Also discussed were Hsp101 (PF3D7_1116800), Exp2 (PF3D7_1471100), and PTEX150 (PF3D7_1436300).

## Acknowledgements

We would like to thank Anna Oksman for technical assistance, Dr. Audrey Odom John (Dept. of Pediatrics, Washington University School of Medicine) for anti-HAD1, Dr. Wandy Beatty (Dept. of Molecular Microbiology, Washington University School of Medicine) for electron microscopy, and Dr. Justin Boddey (Walter and Eliza Hall Institute of Medical Research) for WEHI-842.

## Author Contributions

A.S.N and J.C.N conceived the plasmid platform. A.S.N designed and built pSN054 and the pSN054 construct for PMV with supervision from J.C.N. A.J.P designed and performed PMV experiments under supervision of D.E.G.

## References

1. World Health Organization. 2017. World Malaria Report 2017.

2. Cowman AF, Healer J, Marapana D, Marsh K. 2016. Malaria: Biology and Disease. Cell 167:610–624.

3. de Koning-Ward TF, Dixon MWA, Tilley L, Gilson PR. 2016. Plasmodium species: master renovators of their host cells. Nat Rev Microbiol 14:494–507.

4. Spillman NJ, Beck JR, Goldberg DE. 2015. Protein Export into Malaria Parasite–Infected Erythrocytes: Mechanisms and Functional Consequences. Annu Rev Biochem 84:813–841.

5. Marapana DS, Dagley LF, Sandow JJ, Nebl T, Triglia T, Pasternak M, Dickerman BK, Crabb BS, Gilson PR, Webb AI, Boddey JA, Cowman AF. 2018. Plasmepsin V cleaves malaria effector proteins in a distinct endoplasmic reticulum translocation interactome for export to the erythrocyte. Nat Microbiol.

6. Boddey JA, Hodder AN, Günther S, Gilson PR, Patsiouras H, Kapp EA, Pearce JA, de Koning-Ward TF, Simpson RJ, Crabb BS, Cowman AF. 2010. An aspartyl protease directs malaria effector proteins to the host cell. Nature 463:627–31.

7. Russo I, Babbitt S, Muralidharan V, Butler T, Oksman A, Goldberg DE. 2010. Plasmepsin V licenses Plasmodium proteins for export into the host erythrocyte. Nature 463:632–636.

8. Boddey JA, Carvalho TG, Hodder AN, Sargeant TJ, Sleebs BE, Marapana D, Lopaticki S, Nebl T, Cowman AF. 2013. Role of Plasmepsin V in Export of Diverse Protein Families from the Plasmodium falciparum Exportome. Traffic 14:532–550.

9. Hiller NL, Bhattacharjee S, van Ooij C, Liolios K, Harrison T, Lopez-Estraño C, Haldar K, Lopez-Estrano C, Haldar K. 2004. A host-targeting signal in virulence proteins reveals a secretome in malarial infection. Science 306:1934–7.

10. Marti M, Good RT, Rug M, Knuepfer E, Cowman AF. 2004. Targeting malaria virulence and remodeling proteins to the host erythrocyte. Science 306:1930–3.

11. Chang HH, Falick AM, Carlton PM, Sedat JW, DeRisi JL, Marletta MA. 2008. N-terminal processing of proteins exported by malaria parasites. Mol Biochem Parasitol 160:107–115.

12. Boddey JA, Moritz RL, Simpson RJ, Cowman AF. 2009. Role of the Plasmodium export element in trafficking parasite proteins to the infected erythrocyte. Traffic 10:285–299.

13. Bushell E, Gomes AR, Sanderson T, Anar B, Girling G, Herd C, Metcalf T, Modrzynska K, Schwach F, Martin RE, Mather MW, McFadden GI, Parts L, Rutledge GG, Vaidya AB, Wengelnik K, Rayner JC, Billker O. 2017. Functional Profiling of a Plasmodium Genome Reveals an Abundance of Essential Genes. Cell 170:260–272.e8.

14. Boonyalai N, Collins CR, Hackett F, Withers-Martinez C, Blackman MJ. 2018. Function and essentiality of Plasmodium falciparum Plasmepsin V. bioRxiv 404798.

15. Sleebs BE, Lopaticki S, Marapana DS, O’Neill MT, Rajasekaran P, Gazdik M, Günther S, Whitehead LW, Lowes KN, Barfod L, Hviid L, Shaw PJ, Hodder AN, Smith BJ, Cowman AF, Boddey JA. 2014. Inhibition of Plasmepsin V Activity Demonstrates Its Essential Role in Protein Export, PfEMP1 Display, and Survival of Malaria Parasites. PLoS Biol 12:e1001897.

16. Sleebs BE, Gazdik M, O’Neill MT, Rajasekaran P, Lopaticki S, Lackovic K, Lowes K, Smith BJ, Cowman AF, Boddey JA. 2014. Transition State Mimetics of the *Plasmodium* Export Element Are Potent Inhibitors of Plasmepsin V from *P. falciparum* and *P. vivax*. J Med Chem 57:7644–7662.

17. Gambini L, Rizzi L, Pedretti A, Taglialatela-Scafati O, Carucci M, Pancotti A, Galli C, Read M, Giurisato E, Romeo S, Russo I. 2015. Picomolar Inhibition of Plasmepsin V, an Essential Malaria Protease, Achieved Exploiting the Prime Region. PLoS One 10:e0142509.

18. Hodder AN, Sleebs BE, Czabotar PE, Gazdik M, Xu Y, O’Neill MT, Lopaticki S, Nebl T, Triglia T, Smith BJ, Lowes K, Boddey JA, Cowman AF. 2015. Structural basis for plasmepsin V inhibition that blocks export of malaria proteins to human erythrocytes. Nat Struct Mol Biol 22:590–596.

19. Ganesan SM, Falla A, Goldfless SJ, Nasamu AS, Niles JC. 2016. Synthetic RNA–protein modules integrated with native translation mechanisms to control gene expression in malaria parasites. Nat Commun 7:10727.

20. Godiska R, Mead D, Dhodda V, Wu C, Hochstein R, Karsi A, Usdin K, Entezam A, Ravin N. 2010. Linear plasmid vector for cloning of repetitive or unstable sequences in Escherichia coli. Nucleic Acids Res 38:e88–e88.

21. Pfander C, Anar B, Schwach F, Otto TD, Brochet M, Volkmann K, Quail MA, Pain A, Rosen B, Skarnes W, Rayner JC, Billker O. 2011. A scalable pipeline for highly effective genetic modification of a malaria parasite. Nat Methods 8:1078–1082.

22. Collins CR, Das S, Wong EH, Andenmatten N, Stallmach R, Hackett F, Herman J-P, Müller S, Meissner M, Blackman MJ. 2013. Robust inducible Cre recombinase activity in the human malaria parasite Plasmodium falciparum enables efficient gene deletion within a single asexual erythrocytic growth cycle. Mol Microbiol 88:687–701.

23. Wagner JC, Goldfless SJ, Ganesan SM, Lee MCS, Fidock DA, Niles JC. 2013. An integrated strategy for efficient vector construction and multi-gene expression in Plasmodium falciparum. Malar J 12:1–13.

24. Wagner JC, Platt RJ, Goldfless SJ, Zhang F, Niles JC. 2014. Efficient CRISPR-Cas9–mediated genome editing in Plasmodium falciparum. Nat Methods 11:915–8.

25. Spillman NJ, Beck JR, Ganesan SM, Niles JC, Goldberg DE. 2017. The chaperonin TRiC forms an oligomeric complex in the malaria parasite cytosol. Cell Microbiol 19:e12719.

26. Nkrumah LJ, Muhle R a, Moura P a, Ghosh P, Hatfull GF, Jacobs WR, Fidock D a. 2006. Efficient site-specific integration in Plasmodium falciparum chromosomes mediated by mycobacteriophage Bxb1 integrase. Nat Methods 3:615–21.

27. de Koning-Ward TF, Gilson PR, Boddey JA, Rug M, Smith BJ, Papenfuss AT, Sanders PR, Lundie RJ, Maier AG, Cowman AF, Crabb BS. 2009. A newly discovered protein export machine in malaria parasites. Nature 459:945–949.

28. Ho C-M, Beck JR, Lai M, Cui Y, Goldberg DE, Egea PF, Zhou ZH. 2018. Malaria parasite translocon structure and mechanism of effector export. Nature.

29. Beck JR, Muralidharan V, Oksman A, Goldberg DE. 2014. PTEX component HSP101 mediates export of diverse malaria effectors into host erythrocytes. Nature 511:592–595.

30. Elsworth B, Matthews K, Nie CQ, Kalanon M, Charnaud SC, Sanders PR, Chisholm SA, Counihan NA, Shaw PJ, Pino P, Chan J-A, Azevedo MF, Rogerson SJ, Beeson JG, Crabb BS, Gilson PR, de Koning-Ward TF. 2014. PTEX is an essential nexus for protein export in malaria parasites. Nature 511:587–91.

31. Garten M, Nasamu AS, Niles JC, Zimmerberg J, Goldberg DE, Beck JR. 2018. EXP2 is a nutrient-permeable channel in the vacuolar membrane of Plasmodium and is essential for protein export via PTEX. Nat Microbiol.

32. Chaubey S, Grover M, Tatu U. 2014. Endoplasmic reticulum stress triggers gametocytogenesis in the malaria parasite. J Biol Chem 289:16662–16674.

33. Burrows JN, Duparc S, Gutteridge WE, Hooft van Huijsduijnen R, Kaszubska W, Macintyre F, Mazzuri S, Möhrle JJ, Wells TNC. 2017. New developments in anti-malarial target candidate and product profiles. Malar J 16:26.

34. Banerjee R, Liu J, Beatty W, Pelosof L, Klemba M, Goldberg DE. 2002. Four plasmepsins are active in the Plasmodium falciparum food vacuole, including a protease with an active-site histidine. Proc Natl Acad Sci U S A 99:990–995.

35. Guggisberg AM, Park J, Edwards RL, Kelly ML, Hodge DM, Tolia NH, Odom AR. 2014. A sugar phosphatase regulates the methylerythritol phosphate (MEP) pathway in malaria parasites. Nat Commun 5:4467.

